# Deconfounded Dimension Reduction via Partial Embeddings

**DOI:** 10.1101/2023.01.10.523448

**Authors:** Andrew A. Chen, Kelly Clark, Blake Dewey, Anna DuVal, Nicole Pellegrini, Govind Nair, Youmna Jalkh, Samar Khalil, Jon Zurawski, Peter Calabresi, Daniel Reich, Rohit Bakshi, Haochang Shou, Russell T. Shinohara, the Alzheimer’s Disease Neuroimaging Initiative, the North American Imaging in Multiple Sclerosis Cooperative

## Abstract

Dimension reduction tools preserving similarity and graph structure such as *t*-SNE and UMAP can capture complex biological patterns in high-dimensional data. However, these tools typically are not designed to separate effects of interest from unwanted effects due to confounders. We introduce the partial embedding (PARE) framework, which enables removal of confounders from any distance-based dimension reduction method. We then develop partial *t*-SNE and partial UMAP and apply these methods to genomic and neuroimaging data. Our results show that the PARE framework can remove batch effects in single-cell sequencing data as well as separate clinical and technical variability in neuroimaging measures. We demonstrate that the PARE framework extends dimension reduction methods to highlight biological patterns of interest while effectively removing confounding effects.

## 1 Introduction

Dimension reduction tools such as principal coordinates analysis (PCoA), *t*-distributed stochastic neighbor embedding (*t*-SNE), and uniform manifold approximation and projection (UMAP) are widely employed for exploration of high-dimensional data. These methods all identify lower-dimensional embeddings in Euclidean space that preserve information in the original space. These methods has been demonstrated to reveal complex patterns including cell lineages in single-cell RNA sequencing (scRNA-seq) data (**?**) and neurodevelopmental changes in brain volumetric data (**?**). However, in their current form, these methods do not account for covariates and are known to be substantially influenced by confounders such as batch (Hicks et al., 2018).

Researchers have developed several extensions of dimension reduction tools that are designed for removal of confounding effects. For principal component analysis (PCA), researchers developed PCA with adjustment for confounding variation (**?**). Adjusted PCoA (aPCoA) examines residuals from a linear model on principal coordinates, which are orthogonal to specified confounding variables (Shi et al., 2020). Projected *t*-SNE orthogonalizes the embeddings at each iteration of the *t*-SNE optimization to adjust for batch effects (Aliverti et al., 2020). Another method addresses batch effects by using *t*-SNE to construct a reference embedding based on one batch and then projects observations from other batches onto the reference (Poličar et al., 2021). To date, adjustment for confounders in distance-based dimension reduction methods has required modification of each framework to address this specific problem. Furthermore, many methods including UMAP have not been extended to address confounding.

We develop the partial embedding (PARE) as a generalizable framework for removing nuisance effects from any distance-based dimension reduction method. We achieve this by using the covariate-adjusted dissimilarities from aPCoA as inputs into dimension reduction methods. When the original distances are Euclidean, we can achieve identical results by treating adjusted principal coordinates as input data (see Methods). We refer to these covariate-adjusted dimension reduction results as partial embeddings (PAREs). These PAREs preserve pairwise distances from the original space while removing confounding effects. PAREs can be produced from a broad class of dimension reduction methods including *t*-SNE (van der Maaten and Hinton, 2008), UMAP (McInnes et al., 2020), Laplacian Eigenmaps (Belkin and Niyogi, 2003), diffusion map embeddings (Coifman et al., 2005), LargeVis (Tang et al., 2016), TriMap (Amid and Warmuth, 2022), ForceAtlas2 (Jacomy et al., 2014) and others. Specifically, we apply the PARE framework to *t*-SNE and UMAP to develop partial *t*-SNE (p-*t*-SNE) and partial UMAP (p-UMAP).

## 2 Methods

### 2.1 Adjusted principal coordinates analysis

Let *y_1_, y_2_,…,y_n_* be multivariate observations from samples *i* = 1,2,…,*n*, which can be features from genomics, neuroimaging, or any type of multivariate data. Let *D* = (*d_ij_*)_*n×n*_ denote the sample dissimilarity matrix computed on these observations *y_i_*, where *d_ij_* = *d*(*y_i_*,*y_j_*) and *d* is a chosen dissimilarity function. Define the doubly-centered dissimilarity matrix *G* = (*I* — **11**^*T*^)*A*(*I* — **11**^*T*^) where 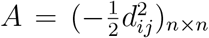. Principal coordinates analysis (PCoA) finds coordinates in Euclidean space that optimally preserve dissimiliarites from the original space. The classical solution finds these coordinates via eigendecomposition of *G* (Gower, 1966). Decomposing *G* = *U*Λ*U*^T^, these coordinates are given by *Z* = *U*Λ^1/2^. Under Euclidean dissimilarities, the principal coordinates *Z* preserve the exact distances from the original space. If the original dissimilarities are non-Euclidean, *Z* may contain imaginary coordinates. Adding a constant to every pairwise dissimilarity can produce coordinates in Euclidean space (Cailliez, 1983).

In adjusted principal coordinates analysis (aPCoA), a linear model is used to remove the effect of nuisance covariates from the principal coordinates (Shi et al., 2020). Let *X* be an *n* × *p* design matrix of nuisance covariates with corresponding projection matrix *H* = *X*(*X^T^X*)^-1^*X^T^*. Covariate-adjusted coordinates are given by *E* = (*I* — *H*)*Z* with adjusted dissimilarity matrix Δ = *EE^T^* = (*I* — *H*)*G*(*I* — *H*). The first two adjusted coordinates are typically used for visualization.

### 2.2 Partial embeddings

We develop the partial embedding (PARE) framework by leveraging aPCoA to remove the effect of confounders from pairwise dissimilarities in the original space. The covariate-adjusted dissimilarity matrix Δ = (*δ_ij_*)_*n×n*_ = (*I* − *H*)*G*(*I* – *H*) is used as an input into dimension reduction methods. For example, UMAP defines affinities based on dissimilarity metrics *d* as *v_ij_* = *v_j_*|*i* + *v_i_*|*j* – *v_j_*|_*i*_*v*_*i*_|*j*

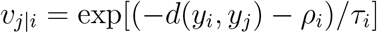

where and *ρ_i_* are the dissimilarity to the nearest neighbor of *y_i_* and *τ_i_* are normalizing factors computed based on dissimilarities among a chosen number of nearest neighbors. A PARE for UMAP can be formulated via the adjusted affinities

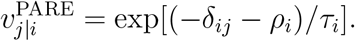

For Euclidean distances, dimension reduction methods can instead take the principal coordinates as inputs. As examples, we highlight how *t*-SNE and UMAP can be equivalently formulated in terms of principal coordinates. *t*-SNE measures similarity in the original space as affinities 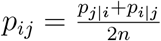 under a Gaussian kernel where

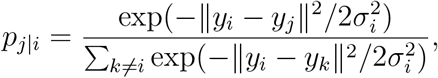

||·|| is the Euclidean norm, and *σ_i_* are chosen to yield a specified perplexity value for each observation. UMAP using Euclidean distances defines similarities using a locally adaptive exponential kernel as *v_ij_* = *V_j_*|*i* + *v_i_*|*j* — *V_j_*|_*i*_*V*_*i*_|*j* where

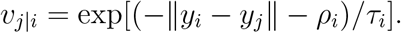

Let *z_i_* denote the principal coordinate vector for observation *i.* For Euclidean distances in the original space, the principal coordinates have identical pairwise distances such that ||*z_i_* − *Z_j_*|| = ||*y_i_* − *y_j_*||. Then the *t*-SNE and UMAP affinities can be written in terms of principal coordinates as

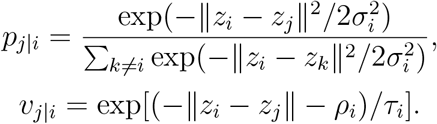

We develop PAREs for any dimension reduction method based on Euclidean distances by instead taking adjusted principal coordinates *e_i_* = (*I* — *H*)*z_i_* as input data. These adjusted coordinates preserve dissimilarities while removing unwanted effects due to the nuisance covariates *X*. We outline the steps in constructing PAREs using Euclidean distances below:

1. Obtain principal coordinates *Z* from the original data from the Euclidean distance matrix *D* as described in subsection 2.1.
2. Using a linear model, residualize *Z* with respect to nuisance covariates *X* to obtain adjusted coordinates *E* = (*I* — *H*)*Z*, where *H* = *X*(*X^T^X*)^-1^*X^T^*.
3. Input the adjusted coordinates *E* to any dimension reduction method based on Euclidean distances.

Obtaining these adjusted coordinates only requires eigendecomposition of the original dissimilarity matrix followed by residualization using a linear model. Both steps are implemented via multiple packages in R, Python, MATLAB, and other programming languages.

For our investigation, we apply our PARE framework to *t*-SNE and UMAP using Euclidean distances to develop p-*t*-SNE and p-UMAP. We use R (version 4.1.1) implementations for *t*-SNE and UMAP in the packages Rtsne (version 0.15) and umap (version 0.2.7.0). Throughout our applications, we choose the perplexity as 10 for *t*-SNE and the number of nearest neighbors as 15 for UMAP.

### 2.3 Human pancreatic cell scRNA-seq data

We apply PAREs to human pancreatic cell scRNA-seq data to remove batch and donor effects from data collected across four separate studies with varying number of cells and RNA-seq protocol. We include RNA-seq data from Baron et al. (2016) (8569 cells, inDrop protocol), Lawlor et al. (2017) (1050 cells, SMARTer), Muraro et al. (2016) (2122 cells, CEL-Seq2), and Segerstolpe et al. (2016) (2133 cells, SMART-Seq2). We treat each study as a separate batch and treat each donor as distinct across studies. We follow a pre-processing pipeline proposed in Lun et al. (2016). First, we use Scran (release 3.15) in R to perform lognormalization and selection of highly variable genes (HVGs) using the counts data from each study. Genes that are not present in all four studies were removed from further evaluation. We then perform normalization by computing size factors across pools of cells, then obtaining factors for each cell via a deconvolution approach (Lun et al., 2016). Within each study, locally weighted scatterplot smoothing (LOESS) is applied to model the mean-variance relationship among genes. We then use a weighted arithmetic mean of mean and variance statistics across studies to select 2,000 HVGs.

We remove cells labelled as “unclear”, “none”, “unclassified” or “co-expression”. After preprocessing, our human pancreatic cell dataset is comprised of 13,369 cells with four batches, 26 donors, and 13 cell types. We apply p-*t*-SNE and p-UMAP to remove batch effect or donor effects in the embeddings. For comparison, we also apply projected *t*-SNE for batch correction (BC-*t*-SNE, Aliverti et al., 2020) with perplexity of 10. We compare our methods visually and numerically using the local inverse Simpson’s index (LISI, Korsunsky et al., 2019). For measuring integration of cells across batches, we compute LISI for batch (bLISI), which captures the effective number of batches in a local neighborhood around each cell. We also examine LISI computed for cell type (cLISI), which captures the number of neighboring cell types and decreases as the separation between cell types increases. We compute bLISI and cLISI across a range of perplexity values, which capture different neighborhood sizes.

### 2.4 ADNI cortical thickness dataset

We apply PAREs to brain cortical thickness data from the Alzheimer’s Disease Neuroimaging Initiative (ADNI) to distinguish technical and biological variability. The data for this study consist of baseline scans which are processed using the ANTs longitudinal single-subject template pipeline (Tustison et al., 2019) with code available on GitHub (https://github.com/ntustison/CrossLong). The ADNI study obtained informed consent from all participants. Institutional review boards approved the study at all of the contributing institutions. Further details for this preprocessing pipeline can be found in Beer et al. (2020).

The full sample consists of 505 subjects, 213 of whom are imaged on scanners manufactured by Siemens, 70 by Philips, and 222 by GE. The sample has a mean age of 75.3 (SD 6.70) and is comprised of 278 (55%) males, 115 (22.8%) Alzheimer’s disease (AD) patients, 239 (47.3%) late mild cognitive impairment (LMCI), and 151 (29.9%) cognitively normal (CN) individuals. We apply p-*t*-SNE and p-UMAP to separate effects of diagnosis and scanner.

### 2.5 NAIMS traveling subjects study

To examine if PAREs can identify technical variability not visible in *t*-SNE and UMAP embeddings, we apply our PAREs to a study of patients with multiple sclerosis (MS) with multiple scan-rescan images across four different sites in the North American Imaging in Multiple Sclerosis (NAIMS) Cooperative. These sites include the University of Pennsylvania (Penn), the Brigham and Women’s Hospital (BWH), the National Institutes of Health (NIH), and the Johns Hopkins University (Hopkins). Nine of the eleven participants are scanned at all four study centers. The mean age of our 11 participants (4 male, 7 female) at time of enrollment was 38 (range 29-47). We received informed consent from all participants, which was approved by the University of Pennsylvania’s institutional review board (IRB).

A standardized high-resoution 3-tesla (3T) MRI brain scan protocol developed by the NAIMS Cooperative was performed at each site (Wattjes et al., 2021). Images were acquired on Siemens Skyra (BWH, NIH), Siemens Prisma (Penn), and Philips Achieva (Hopkins) scanners. Each participant had two scans acquired on the same day at each visit to the study center.

Prior to automated segmentation, images undergo bias correction via nonuniform intensity normalization (N4ITK, Tustison et al., 2010) and FLAIR images are rigidly aligned to the corresponding T1-weighted image within a given scan session. Brain extraction is performed using Multi-Atlas Skull Stripping (MASS, Doshi et al., 2013) and intensity normalization is performed using WhiteStripe (Shinohara et al., 2014). White matter and gray matter volumes are estimated using Joint Label Fusion (JLF, Wang and Yushkevich, 2013), a segmentation method that leverages information from several atlases via weighted voting. These JLF volumes are used as inputs into *t*-SNE, UMAP, and our PARE methods. We use PAREs to identify scanner effects independently of within-subject similarities.

## 3 Results

### 3.1 Case study 1: Human pancreatic cells

We first apply PAREs to analyze scRNA-seq data from human pancreatic cells across four published studies, treated as separate batches (Baron et al., 2016; Lawlor et al., 2017; Muraro et al., 2016; Segerstolpe et al., 2016). We observe clear batch effects in the original *t*-SNE and UMAP visualizations along with a lack of integration among several cell types (Fig. 1). Applying p-*t*-SNE and p-UMAP to remove batch effects considerably reduces separation by batch both visually and numerically, as measured by increases in the local inverse Simpson’s index for batch (bLISI, Supplementary Fig. 1). Remaining batch differences can partially be explained by donor effects, since PAREs with respect to donor show greater visual integration across batches and higher bLISI. PAREs also achieve greater distinction of cell types as measured by decreases in cell type LISI (cLISI, Supplementary Fig. 1). Comparing p-*t*-SNE to the existing projected *t*-SNE for batch correction (BC-*t*-SNE, Aliverti et al., 2020), we find that BC-*t*-SNE achieves greater batch integration but obscures important biological patterns that separate cell types (median cLISI increases from 1.22 in *t*-SNE to 1.32 in BC-*t*-SNE). We also show in **Supplementary Fig. 2** that effective results can be achieved by computing a subset of principal coordinates, which is less computationally intensive. Using scRNA-seq data, we demonstrate that PAREs can uniquely isolate biological variability from unwanted sampling effects in scRNA-seq data.

**Figure 1:**
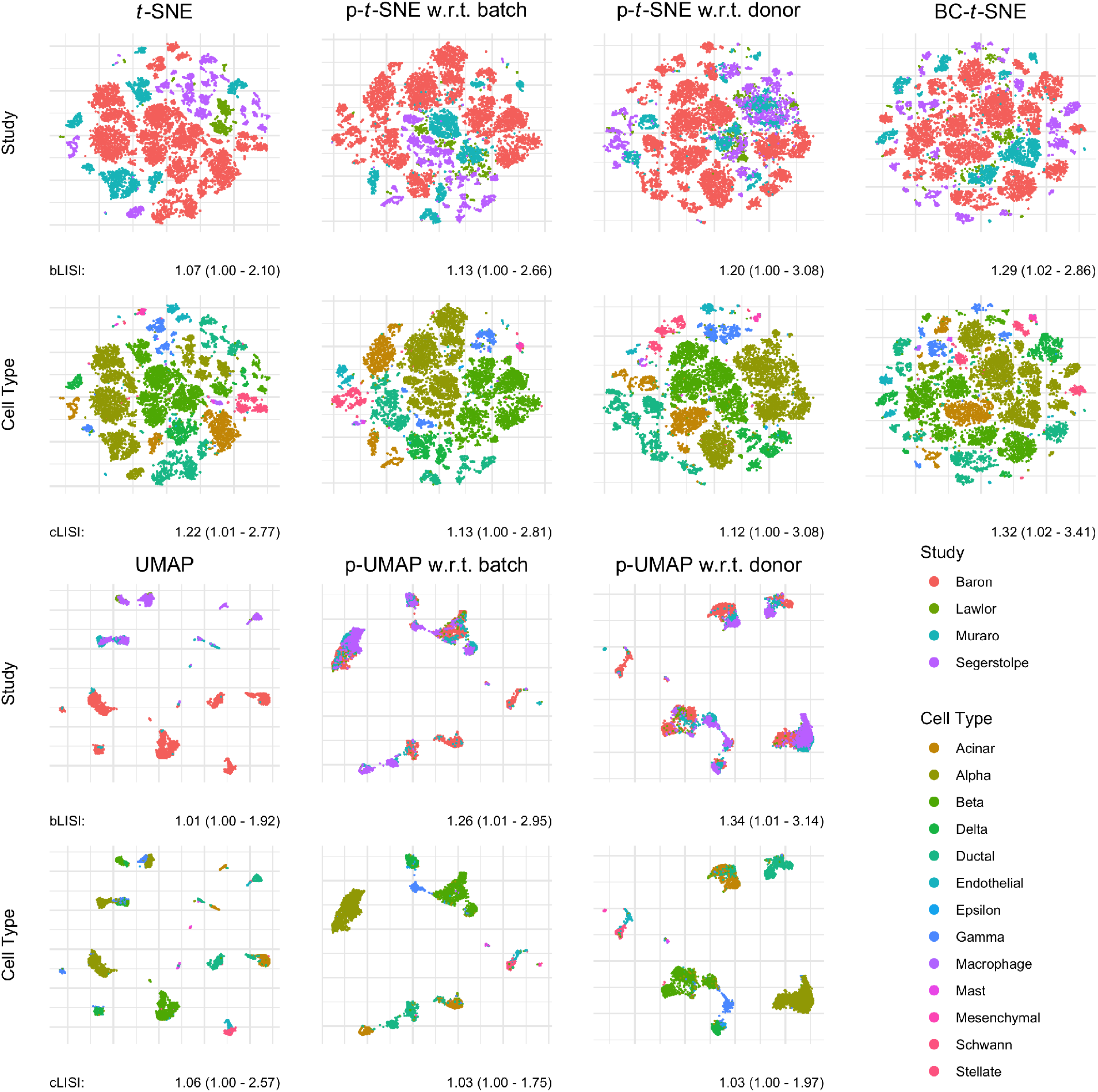
Embeddings and partial embeddings of single-cell RNA-sequencing measurements from 13,369 human pancreatic cells across four studies. The original counts data is log-normalized and reduced to 2,000 highly variable genes. Local Simpson’s index is computed for each cell for batch (bLISI) and cell type (cLISI) with the median, 2.5% quantile, and 97.5% quantile shown. Higher bLISI indicates greater integration across batches and lower cLISI indicates greater separation between cell types. Partial *t*-SNE (p-*t*-SNE) and partial UMAP (p-UMAP) adjust for either batch or donor effects. We compare our new methodology to the existing projected *t*-SNE for batch correction (BC-*t*-SNE). All *t*-SNE embeddings have a perplexity of 10 and UMAP embeddings use 15 nearest neighbors.

**Figure 2:**
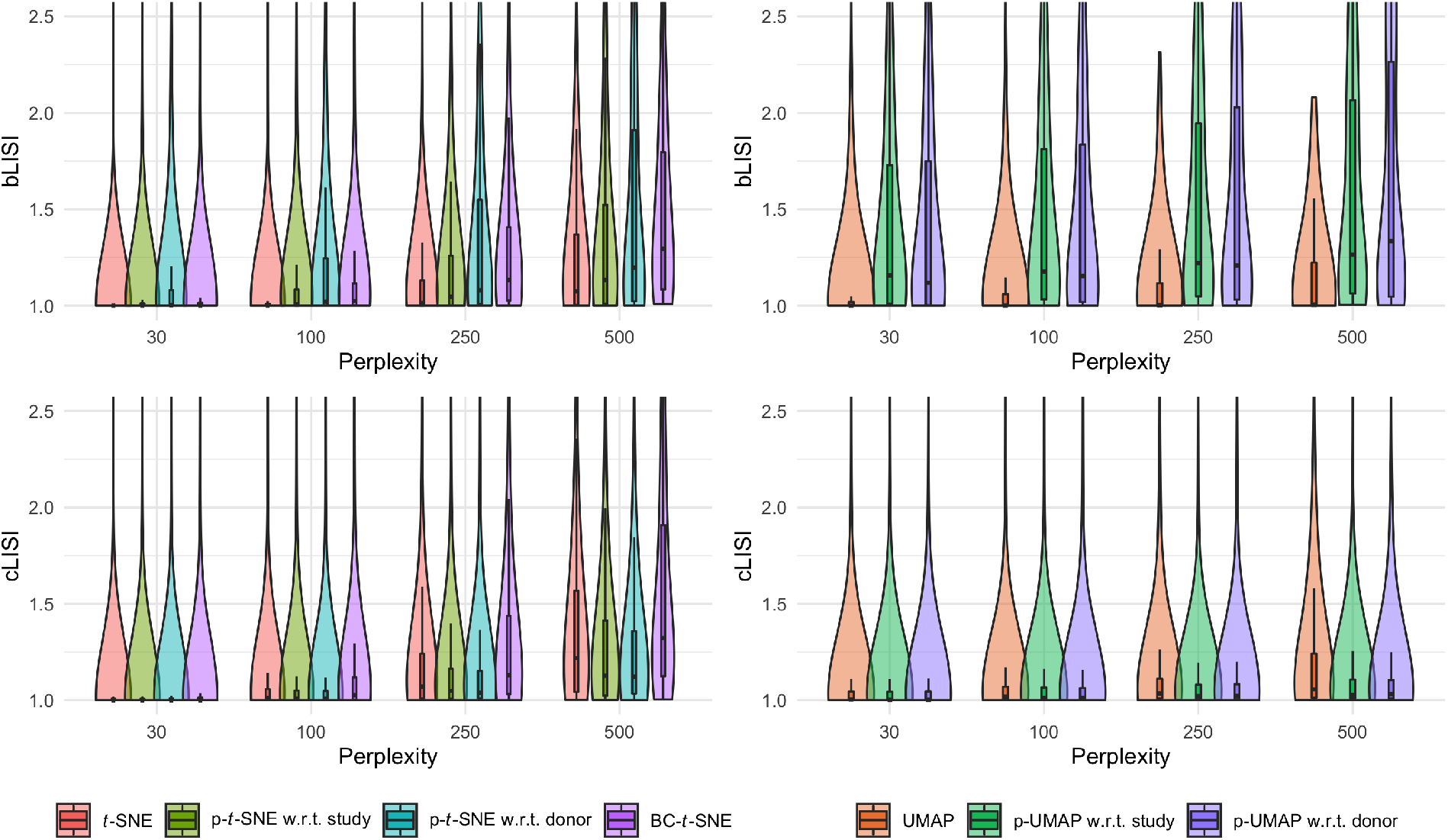
Local Simpson’s index for batch (bLISI) and cell type (cLISI) across multiple perplexity values. LISI is computed using distances from the embeddings. The original embeddings and partial embeddings are compared across perplexity values, which capture different neighborhood sizes around each cell.

**Figure 3:**
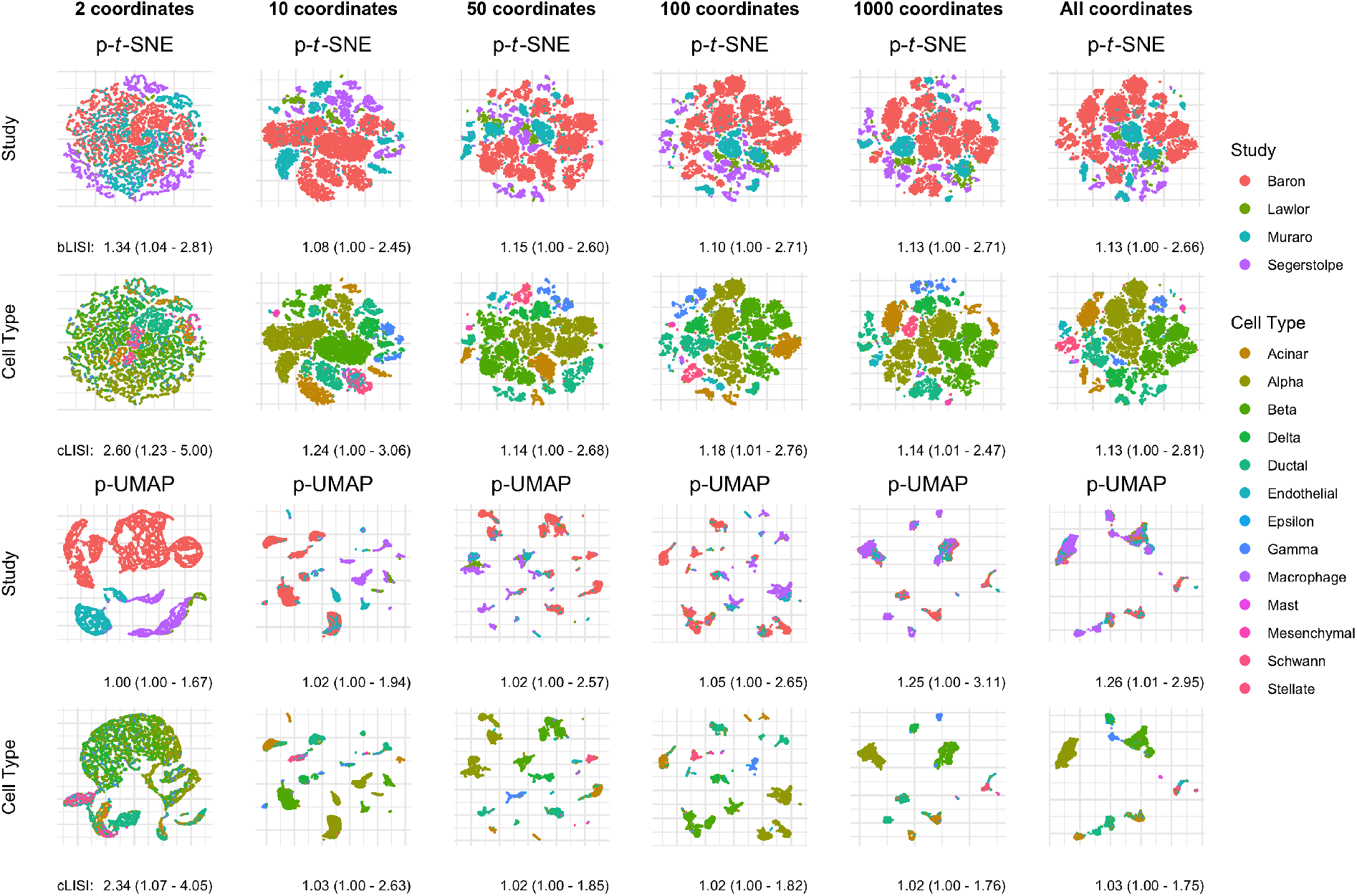
Single-cell RNA-sequencing data embeddings and partial embeddings with respect to batch across varying numbers of principal coordinates. Each dimension reduction method takes a subset of adjusted or unadjusted principal coordinates. The dimension of this subset is varied across figure columns. Partial *t*-SNE (p-*t*-SNE) and partial UMAP (p-UMAP) adjust for batches.

### 3.2 Case study 2: Brain cortical thickness

We next apply PAREs to brain cortical structure measurements to separate biological effects from scanner-related artifacts in the Alzheimer’s Disease Neuroimaging Initiative (ADNI) study. In the ADNI study, researchers previously identified diagnosis-related atrophy in cortical structure and notable batch effects due to differences in scanner properties across study sites (Querbes et al., 2009; Beer et al., 2020). In Fig. 4a, we observe that the original embeddings display both of these effects. However, the confounding scanner effects result in the overlap among images acquired on a Siemens scanner and those from patients with an Alzheimer’s disease (AD) diagnosis. To specifically investigate differences between people with and without AD, we demonstrate that PAREs adjusted for scanner manufacturer maintain diagnosis effects while obscuring scanner influence (Fig. 4a). PAREs are also used to examine scanner effects without the influence of diagnosis effects, highlighting known differences among scanners in the ADNI study.

**Figure 4:**
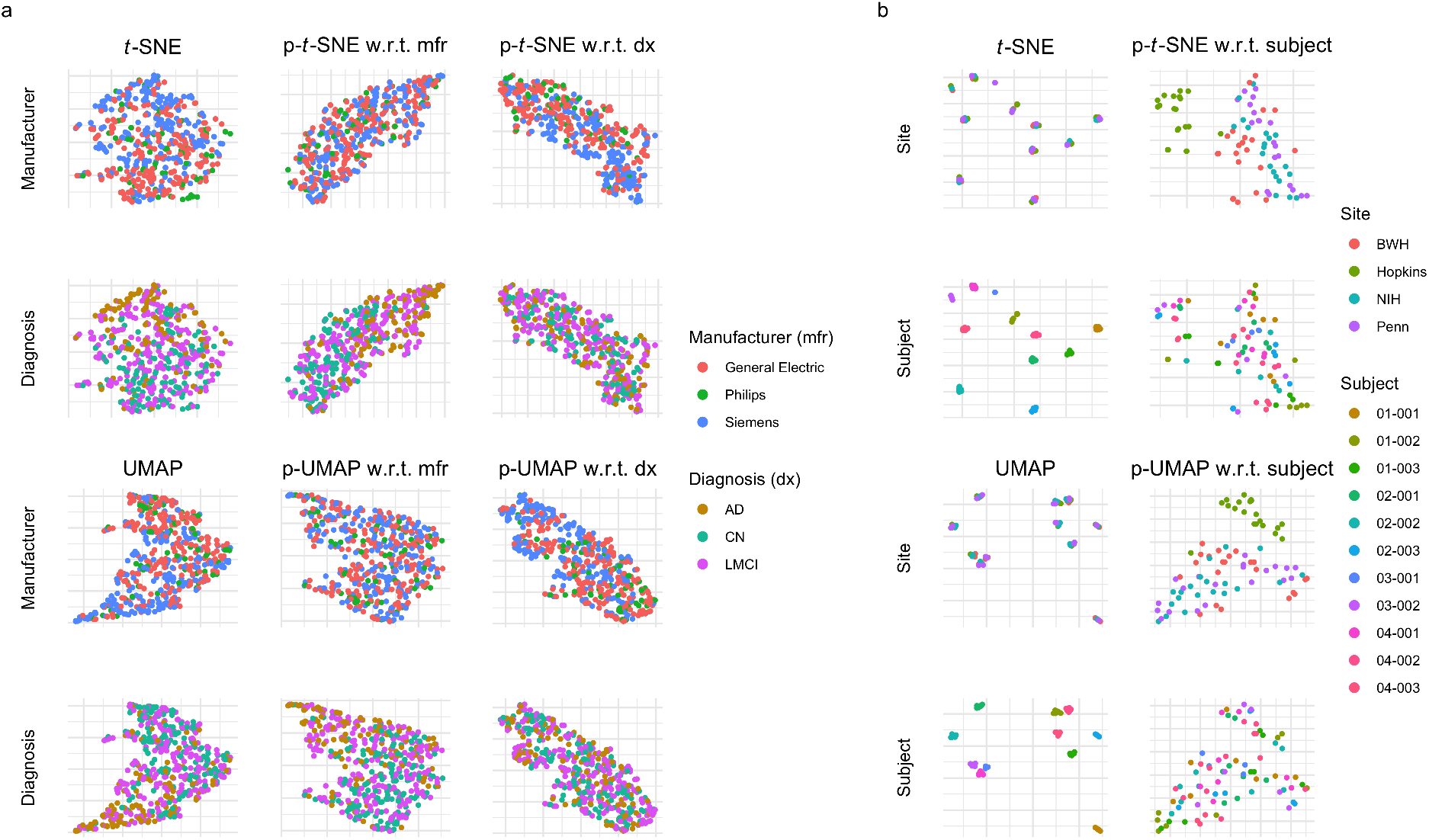
Application of partial embeddings to brain cortical thickness measurements (a) and regional volumes (b) across two neuroimaging studies. **(a)** visualizes cortical thickness data from the Alzheimer’s Disease Neuroimaging Initiative, from which we include 505 participants. These participants are diagnosed as cognitively normal (CN), having late mild cognitive impairment (LMCI), or having Alzheimer’s disease (AD). Participants are acquired across many scanners with three distinct manufacturers. **(b)** shows results from a traveling subjects study of eleven multiple sclerosis (MS) patients with multiple images across four study sites. The Hopkins site uses a Philips scanner while the three other sites use Siemens scanners.

### 3.3 Case study 3: Traveling subject brain volumetric data

Finally, as a proof of concept, we use PAREs to identify scanner effects in brain white matter and gray matter volumes collected as part of a multi-site traveling subjects study of multiple sclerosis (MS). The study involves eleven MS patients with multiple scans across four major imaging centers. We include Siemens images with distortion correction, which was designed to reduce differences with the Philips scanner at Johns Hopkins University (Hopkins). Original *t*-SNE and UMAP embeddings clearly separate white matter and gray matter volumetric measurements by subject regardless of site (Fig. 4b). However, these original embeddings do not capture other types of variability, including potential site effects. We apply p-*t*-SNE and p-UMAP to remove subject effects and discover deviation of images acquired on the Philips scanner at Hopkins from those acquired on Siemens scanners at other sites (Fig. 4b). Here, we show that PAREs can identify technical variability in neuroimaging measures that could not be detected in the original embeddings.

## 4 Discussion

We propose the PARE framework, which extends any distance-based dimension reduction method to adjust for confounders. Our analyses demonstrate that PAREs can be used to target specific patterns in high-dimensional data by removal of confounders. We demonstrate that our proposed PAREs are able to remove batch effects in scRNA-seq exploration, emphasize diagnosis-related changes in brain cortical structure, and identify scanner effects in brain volumetric measurements. For dimension reduction based on Euclidean distances, our PARE framework relies solely on PCoA and linear regression, which are both widely available and computationally efficient. While we only investigate PAREs built on *t*-SNE and UMAP, this framework can be readily applied to a broad class of dimension reduction methods based on distances from the original space.

The PARE framework opens several new directions for methodological development. Future investigations can examine how the PARE framework performs for extensions of methods not considered in this article. PAREs are constructed from the residuals of a linear model, but other models including linear mixed models and general additive models could also be considered for longitudinal and non-linear effects. Extensions of this framework can readily incorporate multiple complex data types by integrating at the level of principal coordinates. Furthermore, PAREs can be extended to examine data types independently of one another by projection of dissimilarity matrices (Székely and Rizzo, 2014). In summary, the PARE framework is able to remove nuisance effects in any distance-based dimension reduction tool. Our framework enables discovery of notable patterns in complex high-dimensional data and introduces a foundation for future methodological research.

## Acknowledgements

This work was supported by the National Institute of Neurological Disorders and Stroke (grant numbers R01 NS085211 and R01 NS060910 and the Intramural Research Program), the National Multiple Sclerosis Society (RG-1707-28586), the National Institute of Mental Health (R01 MH123550 and R01 MH112274), and a seed grant from the University of Pennsylvania Center for Biomedical Image Computing and Analytics (CBICA). The content is solely the responsibility of the authors and does not necessarily represent the official views of the funding agencies.

RB has received consulting fees from Bristol-Myers Squibb and EMD Serono and research support from Bristol-Myers Squibb, EMD Serono, and Novartis. DSR has received research funding from Abata Therapeutics, Sanofi-Genzyme, and Vertex Pharmaceuticals, all unrelated to the current study. RTS receives consulting income from Octave Bioscience and compensation for scientific reviewing from the American Medical Association.

